# Predator: A novel method for targeted protein degradation

**DOI:** 10.1101/2020.07.31.231787

**Authors:** Chuanyang Liu, Jingyu Kuang, Xinyuan Qiu, Lu Min, Wenying Li, Jiaxin Ma, Lingyun Zhu

## Abstract

Protein expression and degradation are fundamental to cell function and physiological status of organisms. Interfering with protein expression not only provides powerful strategies to analyze the function of proteins but also inspires effective treatment methods for diseases caused by protein dysfunction. Recently, harnessing the power of the ubiquitin-proteasome system for targeted protein degradation (TPD) has become the focus of researches. Over the past two decades, TPD technologies, such as E3 ligase modification, PROTACs, and the Trim-Away method, have successfully re-oriented the ubiquitin-proteasome pathway and thus degraded many pathogenic proteins and even "undruggable" targets. However, A low-cost, convenient, and modularized TPD method is currently not available. Herein, we proposed a synthetic biology TPD method, termed Predator, by integrating the classic function of E3 ligase Trim21 and the expression of a bifunctional fusion protein that links Trim21 and the target protein, which leads to the formation of a ternary complex inside mammalian cells and therefore induce the ubiquitination and subsequent proteasome-dependent degradation of the target protein. We first proved this concept by using nanobody and scFv as the targeting module for the Predator system to degrade free GFP and membrane protein ErbB3, respectively. Then, we give an example of how the engineered Predator system can be developed towards biomedical solutions in the context of diabetes mellitus. Ligands-receptor interaction and adenovirus-mediated gene delivery were introduced to the Predator system, and we found this bifunctional fusion protein, in which glucagon was selected to function as the targeting module, downregulated the endogenous glucagon receptor (GCGR) and attenuated glucagon-stimulated glucose production in primary hepatocytes. Although preliminarily, our results showed that this Predator system is a highly modularized and convenient TPD method with good potential for both fundamental researches and clinical usage.

**Graphic abstract:** 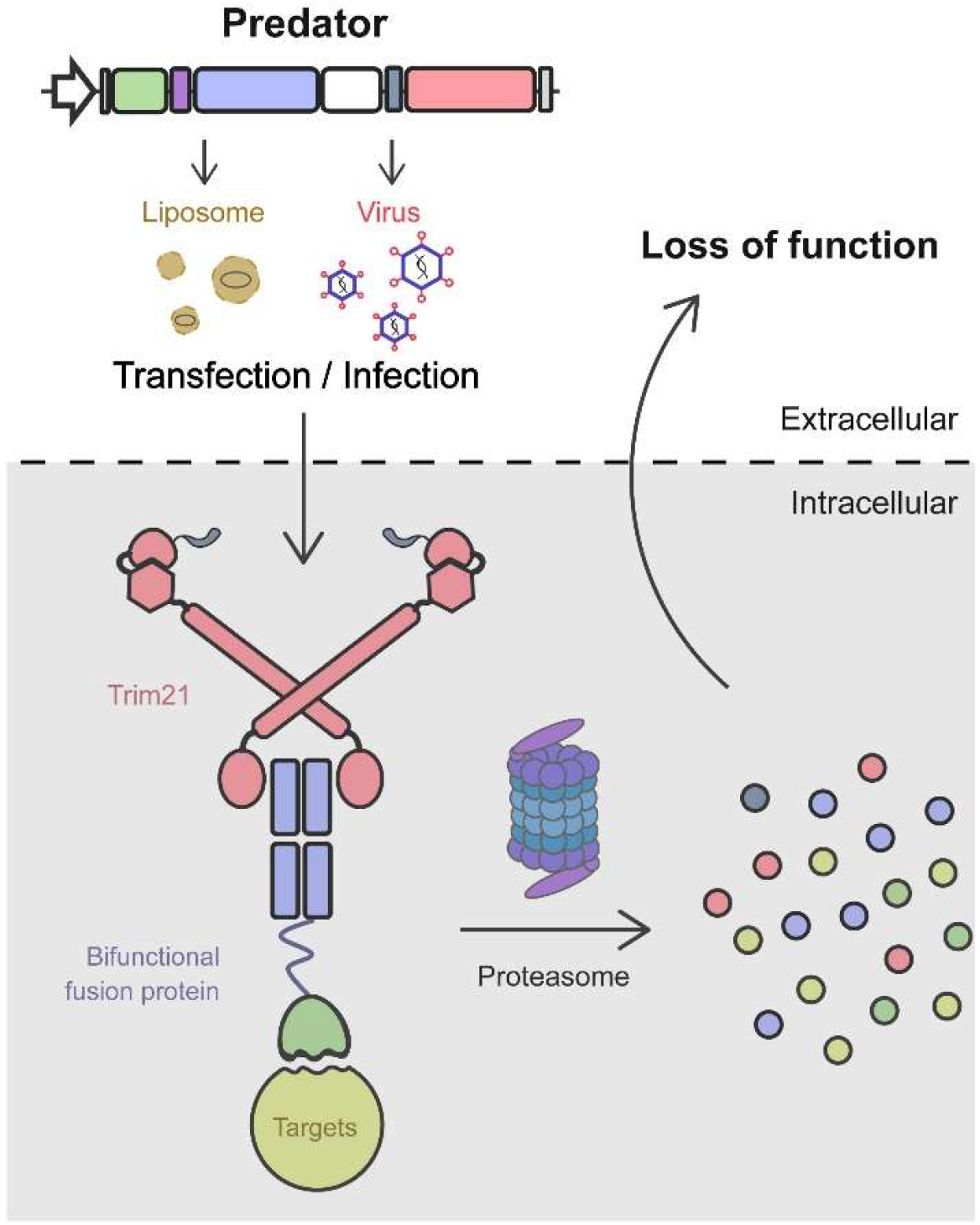

## 1. Introduction

Interfering functions of specific proteins is crucial for both loss-of-function studies and treatment of diseases(*1, 2*). The most commonly used methods are designed to target the production of these target proteins. For example, gene knockout methods(*3–6*) with tools such as CRISPR/Cas9 has been widely used to completely deplete specific protein in host cells. Also, RNA interference(*7*) may lead to the degradation of target mRNA and therefore downregulate the expression of the protein of interest (POI). These methods work on the DNA or RNA level, with significant advantages such as high repression efficiency. However, with these methods, the time required for POI depletion depends on the half-life of the POI in cells, which can be problematic when studying highly dynamic processes, such as the cell cycle, differentiation, or neural activity(*1, 8*). Moreover, these methods were powerless when it comes to long-lived protein targets(*9*). The development of rapid targeted protein degradation tools is hence highlighted recently.

In mammalian cells, most proteins that are destined to be degraded are processed through the ubiquitin-proteasome system (UPS)(*10*). The UPS operates through a collection of regulated and orchestrated steps in which Ub-activating (E1), Ub-conjugating (E2), Ub-ligase (E3), and deubiquitinase (DUBs) enzymes works coordinately, marked target proteins for 26S proteasome degradation by covalent post-translational modification with the ubiquitin(*10, 11*). Within the UPS, E3 ligases, a large gene family with ~600 predicted members, are unique in their role of dictating target specificity(*12*).

Owing to the insights into this high-selectivity and high-efficiency degradation machinery, researchers have developed several tools for the direct degradation of endogenous protein by utilizing the ubiquitin-proteasome system, such as PROTAC, deGradFP, Trim-Away, *etc*.(*13–15*). Trim-Away is a novel strategy through microinjecting Trim21 protein and the antibody of target protein into cells. It harnesses the UPS to remove unmodified native proteins and allows the study of protein function in diverse cell types(*16*). The core fragment of Trim-Away is Trim21 protein, an E3 ubiquitin ligase, that usually presents as a homodimer and binds with high affinity to the Fc domain of antibodies, hence forming a ternary complex consisting of Trim21 antibody-antigen. Then, Trim21 recruits the ubiquitin-proteasome system to antibody-bound antigens, leading to their destruction(*17–19*). This method showed an acute and rapid degradation for target protein exactly at the protein level, but the short lifespan of injected-antibody, the high cost of antibody production, and the technique complexity limit its wide application, especially for clinical usage(*11, 16, 20*).

Here we demonstrate a low-cost, uncomplicated, and high-modularized TPD method, termed Protein Predator, based on the reported biological function of Trim21. The Predator system functions through the expression of Trim21 and a modularized bifunctional fusion protein. The fusion protein is consisted of a targeting module specifically binds to the target protein and recruitment module that recruit Trim21 (Fig 1A). Through liposome or other commonly used gene delivery methods, the Predator system can be introduced into the tissue or cultured cells, and theoretically form a ternary complex with the target protein (Fig 1B), leading to the ubiquitylation of complex and, subsequently, the degradation of the target protein. We believe that the Predator system might add new parts and tools to the synthetic biology toolbox and fundamental researches, expanding the current TPD methods that might foster novel advances in TPD-based therapies.

**Figure 1.**
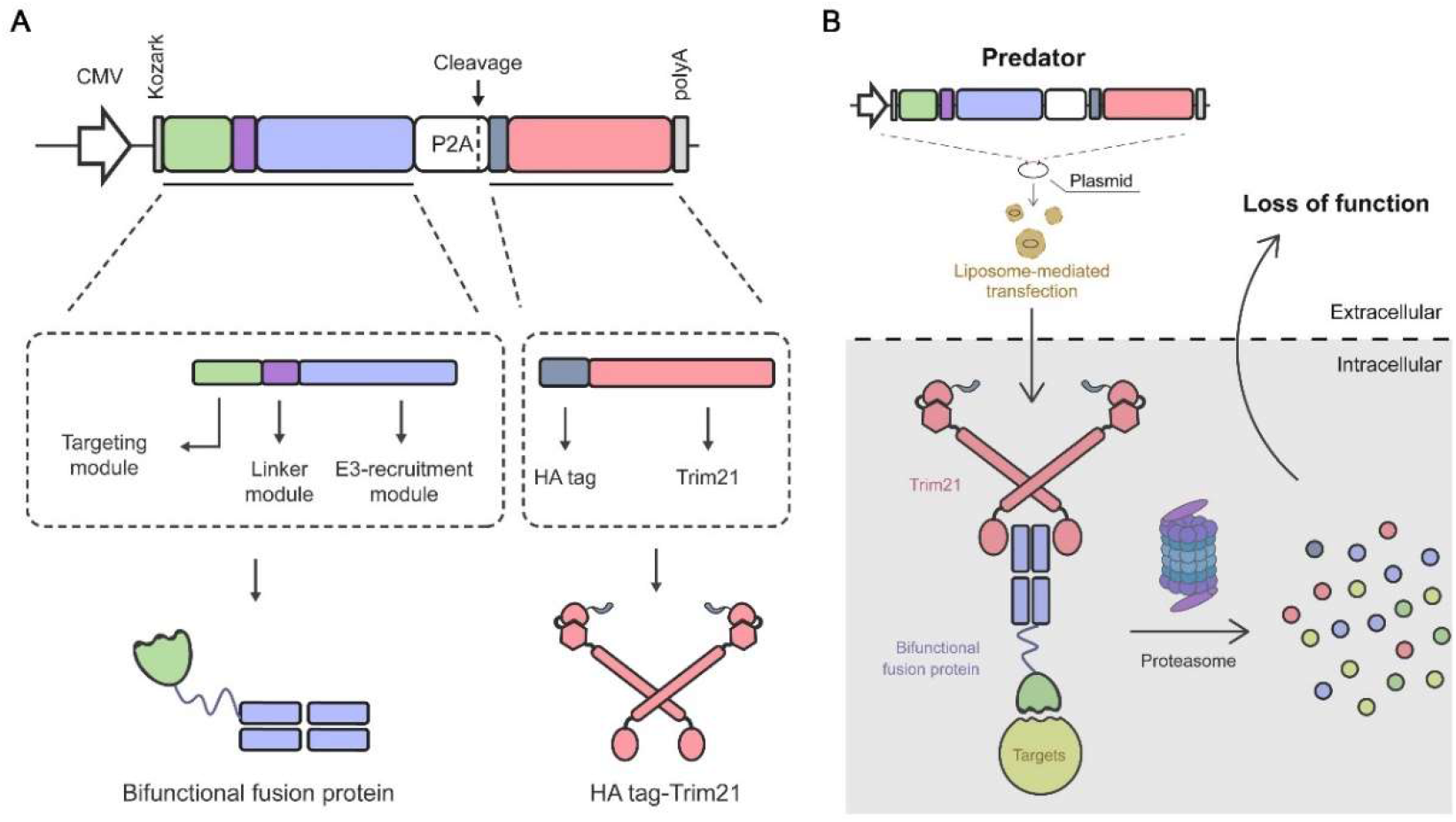
Design of the modularized Predator system.

## 2. Results

### 2.1 Design of Predator system

We attempted to design a novel protein degradation strategy based on the classic features of Trim21 as described. We redesigned a bifunctional fusion protein following a modular design methodology in synthetic biology, which is composed of targeting modules that specifically recognize proteins (e.g., variable region sequences in antibodies that specifically bind antigens) and E3-recruiting module to recruit E3 ligase to the target protein and induce ubiquitination modifications (e.g., IgG conserved region sequence Fc in antibodies that recruit the E3 ligase Trim21 protein), and finally, the linker modules that ensure that these two modules function independently (Fig 1A).

These three modules function together as a substitute for the antibody that links antigen and Trim21 during the degradation of antigen, thereby contribute to the formation of a Trim21-bifunctional fusion protein-target protein ternary complex. Moreover, HA Tag was added to the N-terminal of Trim21, making it easier to be detected. The plasmid which did not express the bifunctional fusion protein and Trim21 was set as the control.

Theoretically, the degradation of target proteins can be achieved by the simultaneous introduction of bifunctional molecular sequences designed for target proteins and Trim21 sequences into the cell, which can be transferred to eukaryotic gene expression vectors via liposomal transfection, cationic polymer transfection techniques which are commonly used in cell biology research. The key components of the degradation system will be automatically generated through gene expression in cells, which, subsequently, leads to the recognition and degradation of the target protein (Fig 1B). This targeted protein degradation system was termed as the Protein Predator.

### 2.2 GFP Predator: based on nanobody targeting module

To verify the degradation efficiency of target protein, we first set up a proof of concept experiment in mammalian cell culture with GFP, a well-established green fluorescent protein used in cell biology research, as the target protein.

For such matters, we first constructed the GFP Predator plasmid expressing following modularized parts: 1) the bifunctional fusion protein, GFP-nanobody (nano)-IgG Fc (IgG Fc), encoding the fusion protein of GFP single domain antibody (nanobody, or nano) and human IgG Fc fragment to target and bind exogenously expressed GFP in the cytoplasm, forming the nano-IgG Fc-GFP complex; 2) HA Tag-Trim21, the PRYSPRY domain of which recognizes the Fc fragment of the nano-IgG Fc -GFP complex and induce the ubiquitination of complex, subsequently leading to the proteasomes-dependent protein degradation (Fig 2A).

**Figure 2.**
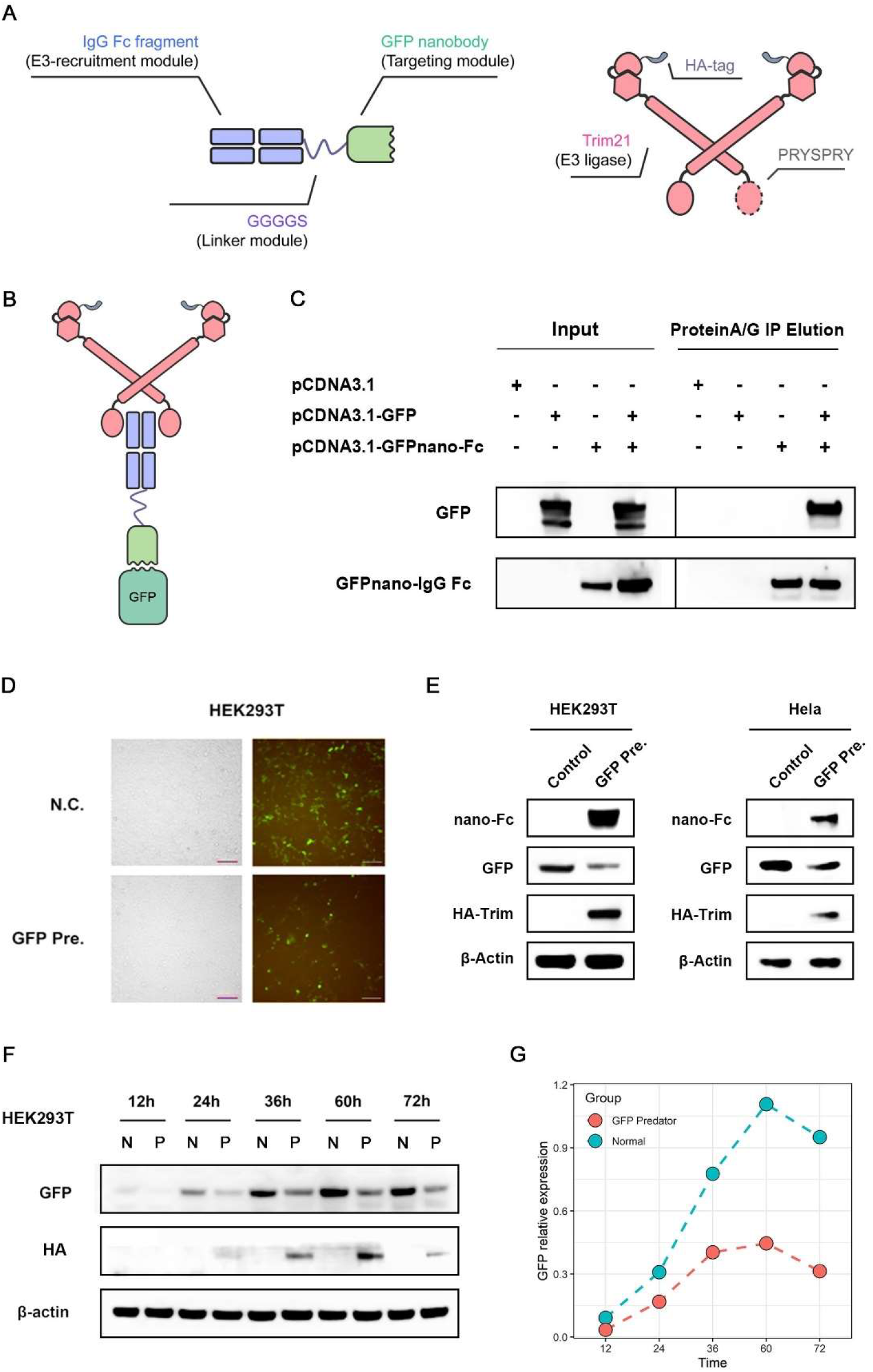
GFP Predator based on nanobody targeting module. Schematic representation showing the modularized design of GFP Predator (GFP Pre.) (A), and its degradation mechanism (B). (C) Immunoprecipitation assay verifying the binding of GFP-nanobody-Fc and GFP. (D) Fluorescence images of the GFP-Predator transfected group and its negative control transfected with an empty vector. HEK293T cells in both groups were transfected with GFP-expression plasmid (E) Western blotting determining the expression level of GFP-nanobody(nano)-IgG Fc (IgG Fc), GFP, HA Tag-Trim21 in HEK293T, and Hela cells transfected with GFP Predator. (F and G) Western blotting assay showing the degradation of GFP in GFP Predator transfected groups at different times after transfection. N, P represents Negative control and GFP Predator, respectively.

To experimentally prove the interaction of the Predator system and target protein, immune precipitation was performed. Western blotting results showed a significant co-precipitation of GFP nanobody and GFP protein (Fig 2B and C), suggesting the interaction of GFP nanobody and GFP protein in HEK293T cells. To test whether the GFP Predator system could induce the degradation of GFP, GFP-expression plasmid and GFP Predator plasmid was co-transfected into HEK293T cells. Fluorescent imaging showed a significant decrease of green fluorescence in GFP Predator transfected groups 48 hours after transfection indicating the decrease of GFP protein level (Fig 2D). The western blotting analysis also showed that GFP was significantly degraded to about 30% of the original level with the appearance of GFP-nano and HA-Trim (Fig 2E), which confirmed that Predator could be used to degrade target protein with high efficiency.

To further reveal the durability of GFP degradation by GFP Predator, we collected cells on 12, 24, 36, 60, and 72 hours post-transfection. As is shown in Figure 2F, western blotting assay revealed that the GFP level of the control group (transfected with GFP-expressing plasmid and Predator control) increased in 12~60 hours, indicating the expression of GFP by GFP-expression plasmid. While the GFP level in cells transfected with Predator was significantly lower than control in each time point. Of note, the degradation efficiency remains increasing even 72h after transfection (53.50% in 60h and 62.38% in 72h), indicating the durability of our Predator (Fig 2G).

### 2.3 ErbB3 predator: using ScFv as the targeting module

To further explore the scalability of the Predator system and its efficiency on proteins which are in different physiological states. The single-chain antibody (ScFv), another class of specific antigen-binding module, was therefore selected for the design of the targeting module of Predator instead of the nanobody. Meanwhile, ErbB3, a transmembrane protein of the epidermal growth factor receptor (EGFR) family, was selected as the target for further exploration of Predator for its special cellular locations, clear downstream functions, and easy-to-measure phenotypes. Hence, we replaced the targeting module of GFP (GFP-nanobody) with the ScFv(*21*) that targets ErbB3(ScFv_ErbB3_, Fig 3A), which transformed GFP Predator into ErbB3 Predator easily.

**Figure 3.**
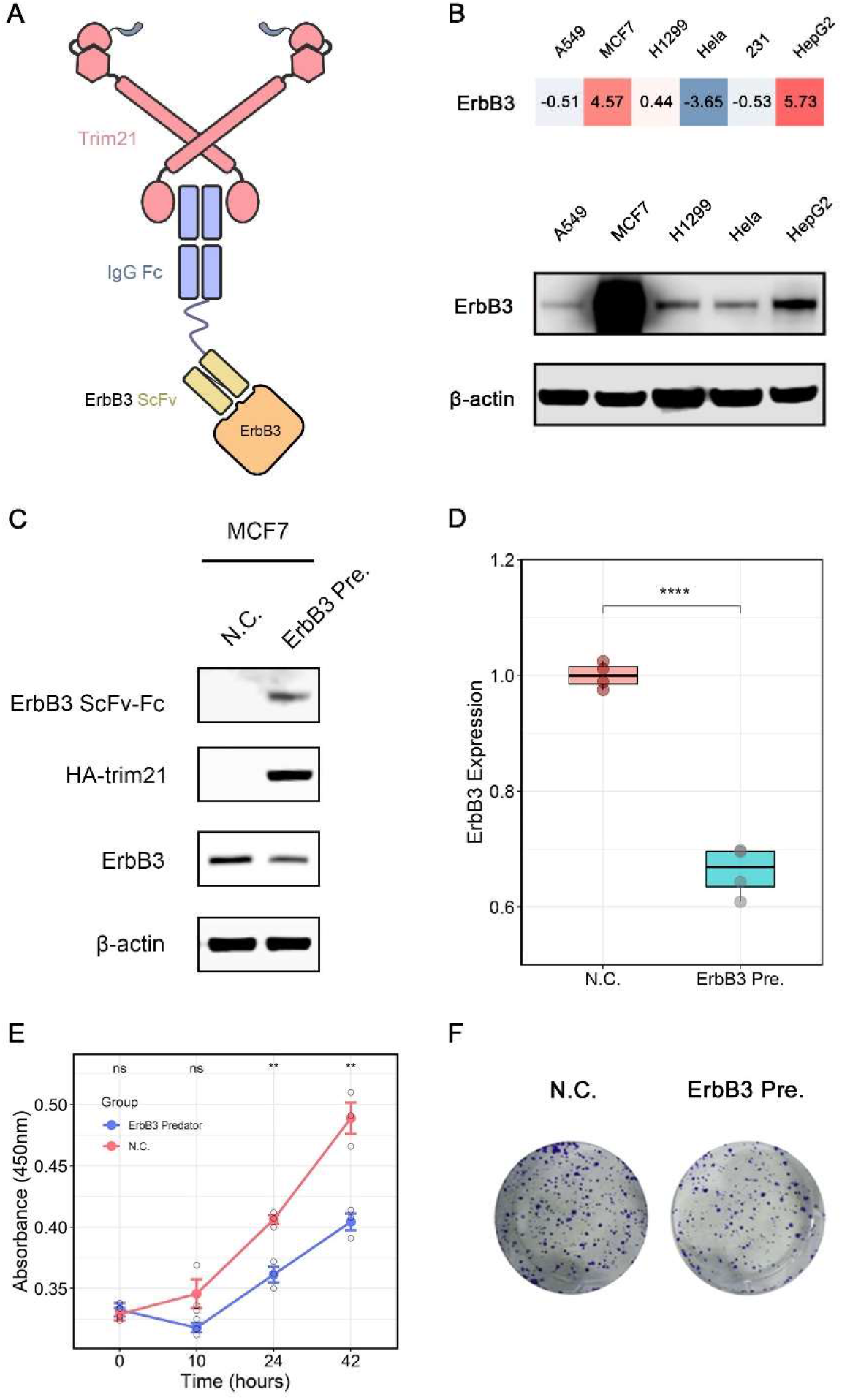
ErbB3 Predator based on the ScFv targeting module. Schematic representation showing the degradation mechanism of ErbB3 Predator (ErBb3 Pre.) (A). (B) the expression level of ErbB3 in cell lines in CCLE database and western blot analysis of ErbB3 expression in commonly-used cell lines. high expression is shown in red. (C and D) Western blotting assay showing the components of ErbB3 Predator and the degradation of ErbB3. CCK8 cell viability assay (E) and clone formation assays (F) showing the proliferative capacity of MCF7 cells. Relative gene expression was calculated using the grayscale of the bands of Western blot. The expression levels of each gene in the control groups were arbitrarily set to 1.0. Error bar represents SD of at least 3 biological replicates, * p<0.05, ** p<0.01.

We first inquired the expression level of ErbB3 in the Cancer Cell Line Encyclopedia (CCLE) database(*22*), which showed that among commonly-used cell lines, MCF7 and HepG2 were high expressing ErbB3. Confirmed to these results, our result from western blot showed that the expression of ErbB3 in MCF7 was highest (Fig 3B).

To validate the protein degradation efficiency of the ScFv-based ErbB3 Predator system, we transfected MCF7 cells with plasmids expressing ScFv_ErbB3_-linker-Fc fusion protein and HA-Trim21. Western Blotting analysis showed that with the high expression of ScFv_ErbB3_-linker-Fc fusion protein and HA-Trim21, the ErbB3 protein level decreased dramatically in cells transfected with ErbB3 Predator (approximately 30%, Fig 3C). This result demonstrated that the ScFv-based Predator system could successfully degrade the targeted membrane protein.

Since it has been established that interfering ErbB3 level significantly suppressed the proliferative capacity of the breast cancer cell lines(*23–26*). We performed CCK8 cell viability assay and clone formation experiments to further demonstrate the effectiveness of Predator-mediated protein degradation of ErbB3. The results showed that the proliferative capacity of MCF7 cells introduced with ErbB3 Predator was significantly weaker than the control cells (p<0.01, Fig 3E), and their clone formation rate were also significantly weaker compared to the control group (Fig 3F).

These results demonstrate that the ErbB3 Predator can inhibit cell proliferative capacity through the degradation of ErbB3 as expected, and also show the great potential of the Predator system for application in fundamental studies.

### 2.4 GCGR predator: from antibody to ligands

Type 2 diabetes (T2D) is the most frequent metabolic disorder around the world(*27*). Glucagon is a 29 amino acid fusion protein hormone secreted by pancreatic alpha cells that has a strong stimulatory effect on hepatic glucose production and acts as a counter-regulatory hormone to insulin to raise blood glucose levels via glucagon receptor (GCGR), a 54kDa G-protein coupled receptor(*28, 29*).

Previous reports showed that liver-specific knockdown of GCGR resulted in decreased fasting glucose, improved insulin sensitivity, and glucose tolerance in both chow diet and high fat diet-fed mice(*30, 31*). These results further suggest that GCGR may serve as a potential target for glycemic control.

To design a Predator system that can specifically degrade the glucagon receptor (GCGR), we replaced the GFP/ErbB3 targeting module with glucagon, which could bind to GCGR specifically. If our Predator system can degrade GCGR protein levels, then the cells would have decreased GCGR protein levels, resulting in decreased sensitivity to glucagon and a corresponding reduction in the activation of glycogenolysis and gluconeogenesis, and eventually glucose production under the stimulation of glucagon in the liver cells (Fig 4A) (*28, 32*).

**Figure 4.**
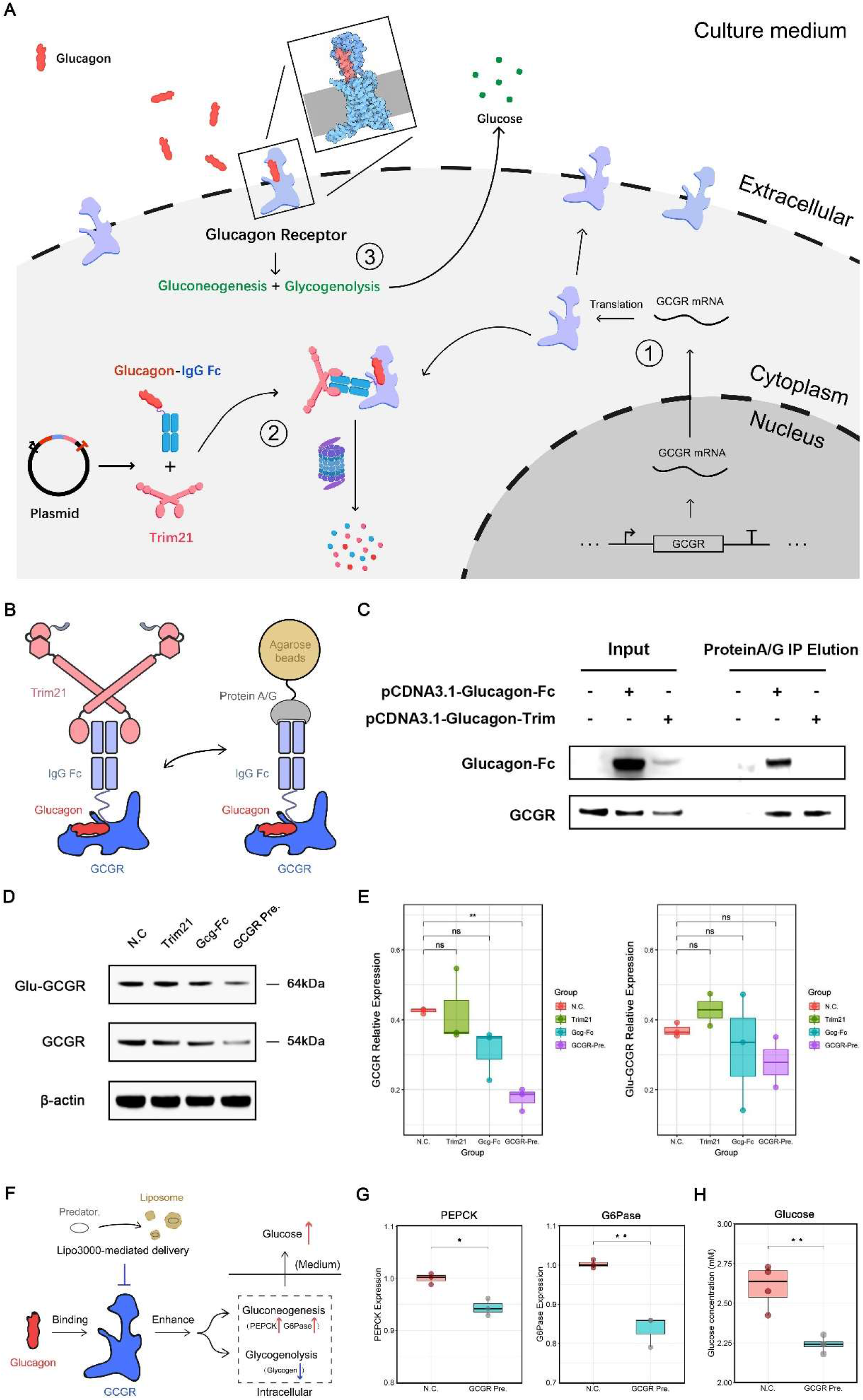
GCGR Predator based on ligands targeting module. (A) Schematic representation showing the mechanism of GCGR Predator. (B and C) Immunoprecipitation assay that verifies the binding of bifunctional fusion protein glucagon-Fc and GCGR. (D) Western blotting assay showing the degradation of GCGR in the GCGR Predator transfected group and its quantified results (E). (F) Schematic representation showing the mechanism that GCGR Predator negatively regulates the glucose metabolic programs of HepG2 cells. (G) Quantitative PCR assay of G6Pase and PEPCK in HepG2 cells transfected with GCGR predator. (H) Glucose production analysis of HepG2 cells transfected with GCGR predator. Relative gene expression was calculated using the 2^−ΔΔCT^ method, with initial normalization of genes against Hprt1 mRNA within each group. The expression levels of each gene in the control groups were arbitrarily set to 1.0. Error bar represents SD of at least 3 biological replicates, * p<0.05, ** p<0.01.

To validate that intracellular glucagon is capable of binding intracellular GCGR, HepG2 cells were transfected with Glucagon-Fc expressing cassette or GCGR Predator (GCGR Pre.) expressing cassette, and immunoprecipitation was performed with IgG-Fc targeting Protein A/G beads 48 h post transcription followed by Western blotting analysis probing GCGR and the Glucagon-Fc domain of the GCGR Predator (Fig 4B). Results showed that interaction signal between Glucagon-Fc and GCGR could be observed on Glucagon-Fc transfected group while the non-transfected group showed no corresponding signal. Surprisingly, cells transfected with GCGR Pre. plasmids showed significantly weaker interaction signals together with a significantly lower GCGR abundance, we reasoned that might due to the Trim21 mediated GCGR degradation (Fig 4C).

We then performed Western Blotting to further prove that such interaction is capable of triggering GCGR degradation. Results indicated that GCGR Pre. transfected cells showed a ~40% lower GCGR level compared to the vector control (Fig 4D and E) and this degradation relies on proteasome (Data not shown). Besides, there is no significant difference between the control group and groups that only overexpressed Glucagon-linker-IgG Fc (Gcg-Fc) or Trim21. Also, we found that both in four groups, Glycosylated GCGR (Glu-GCGR, ~64kDa) showed no difference in expression level.

Since changes in glucose metabolic programs are expected under GCGR depletion(*30, 31*), we performed the metabolic analysis to determine if our GCGR Predator could generate similar phenotype changes. For such assay, HepG2 cells were transfected with GCGR Predator by lipofectamine 3000 and stimulated with Glucagon in DMEM medium without glucose (Fig 4F). To reveal the metabolic changes generated by GCGR Predator, we used quantitative PCR analysis to evaluate the cellular gluconeogenesis process. Results showed that GCGR Predator transfection significantly down-regulates the expression of key enzymes in gluconeogenesis pathway, Glucose 6-phosphatase (G6Pase), and Phosphoenolpyruvate carboxykinase (PEPCK) (Fig 4G). To measure the overall glucagon sensitivity, glucose concentration in the medium was evaluated by glucose oxidase assay. As shown in Figure 2b, GCGR predator carrying cells showed significantly lower glucose concentration (~20% decrease in HepG2, Fig 4H) in the culture medium comparing to the control group, indicating a significantly impaired overall glucagon sensitivity mediated by GCGR Predator. Generally, these results indicated that the GCGR predator significantly impairs hepatocyte glucagon sensitivity *in vitro* by affecting metabolic programs downstream of GCGR.

### 2.5 Adenovirus-based delivery of GCGR Predator to primary cells

Clinical application potential of current protein degradation techniques such as Trim-Away, AID, and deGradFP was thought to be limited by their technique complexity(*8, 10, 11, 16, 20*). To further explore the clinical potential of our Predator system, the GCGR Predator system was constructed into an adenovirus-based gene therapy vector that is widely used in pre-clinical applications, and primary hepatocytes were selected for degradation and downstream phenotypic validation of the GCGR Predator system as it is closer to normal physiological condition.

To design a Predator system that works in mice, we first acquired the mouse homologous gene sequences of Glucagon, IgG Fc, and Trim21 at NCBI and replaced corresponding sequences in the original human GCGR Predator system with this modules, and then inserted the complete mouse GCGR Predator sequence into the adenovirus vector AdTrack and packaged the adenovirus to infect primary hepatocytes (Fig 5A).

**Figure 5.**
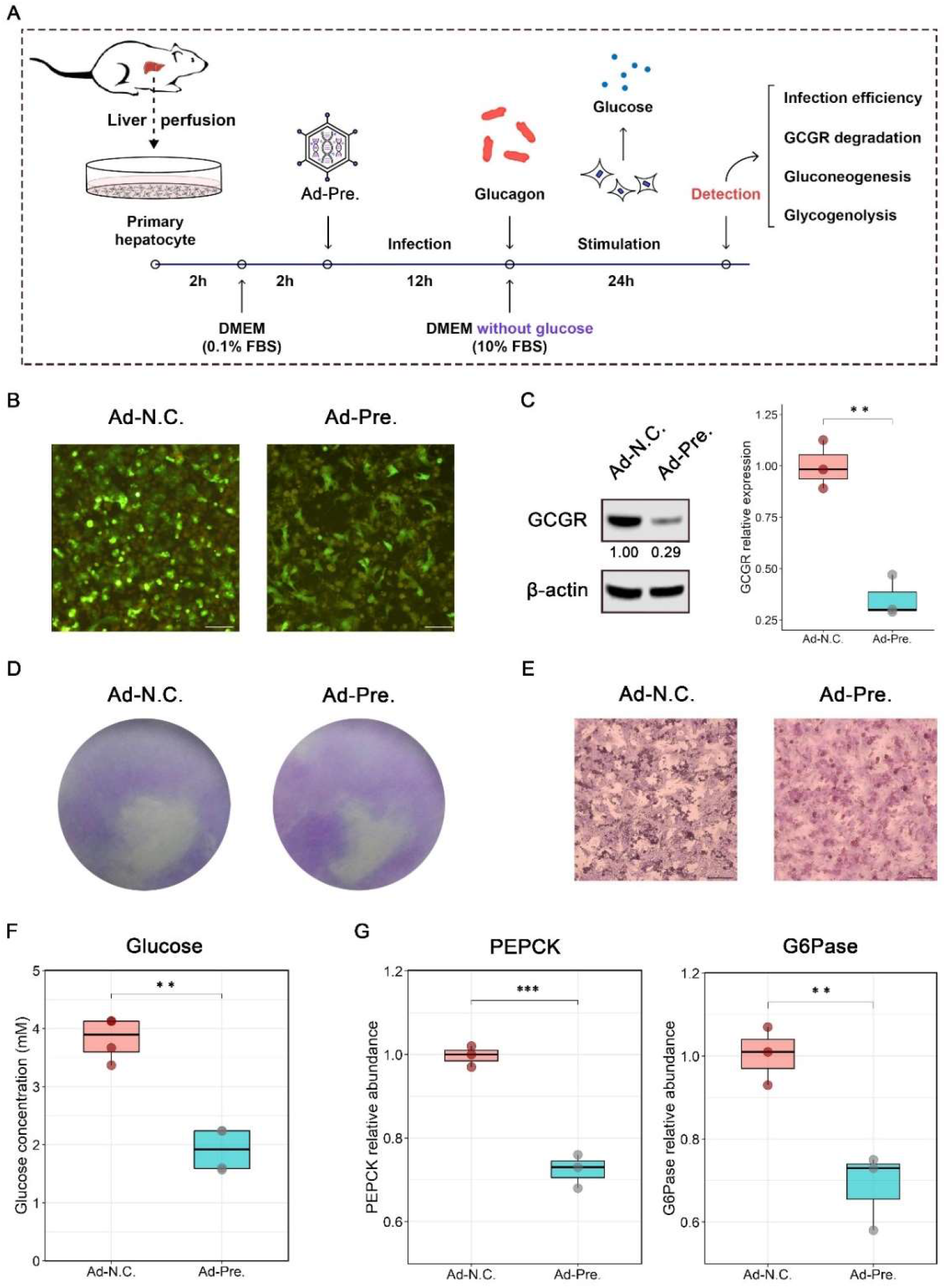
Adenovirus-based gene delivery of GCGR Predator to primary hepatocytes. (A) Schematic representation showing the workflow. (B) Fluorescence images of the Adenovirus-mediated GCGR-Predator (Ad-Pre.) infected group and its negative control Adenovirus (Ad-N.C.) infected with an empty vector Ad-N.C. (C) Western blotting assay showing the degradation of GCGR in the GCGR Predator transfected group (D) Representative glycogen staining microscopy of GCGR predator infected Primary hepatocytes. (F) Glucose production analysis of primary hepatocytes infected with Adenovirus-GCGR Predator. (G) Quantitative PCR assay of G6Pase and PEPCK in primary hepatocytes transfected with GCGR predator. (H) Relative gene expression was calculated using the 2^−ΔΔCT^ method, with initial normalization of genes against mouse Hprt1 mRNA within each group. The expression levels of each gene in the control groups were arbitrarily set to 1.0. Error bar represents SD of at least 3 biological replicates, * p<0.05, ** p<0.01.

After collecting mouse primary hepatocytes via liver perfusion, the primary cells were infected with GFP fluorescence-labeled adenovirus (carrying GCGR Predator or negative control), and the delivery efficiency of the Predator system was judged by measuring the cellular GFP fluorescence after 36 h. The results showed high efficiency of adenovirus infection in both the control and the experimental groups (Fig 5B), which confirmed the delivery of the Predator system to primary cells.

To verify the degradation effect of GCGR Predator on GCGR in primary hepatocytes, Western blot analysis on GCGR was performed in primary hepatocytes infected with either GFP expressing adenovirus or GCGR Predator expressing adenovirus. Results showed a significant down-regulation of GCGR levels in the experimental group of primary cells expressing GCGR Predator (approximately 70% down-regulation, Fig 5C), suggesting that the adenovirus-delivered GCGR Predator system successfully degrades the target protein GCGR in primary hepatocytes.

Similarly, we tested the effect of GCGR Predator on the downstream glycogenolysis process and the gluconeogenesis pathway. The results showed that the glycogenolysis process of GCGR Predator group was significantly inhibited (Figure 5E), and the mRNA levels of the key enzymes PEPCK and G6Pase in its gluconeogenesis pathway were also relatively down-regulated (Figure 5G, p<0.001 for PEPCK and p<0.01 for G6Pase), as well as the overall glucose production (Fig 5F, p<0.001). These results for primary hepatocytes are similar to the results of HepG2, showing that the adenovirus-based gene therapy vector can successfully achieve the delivery of GCGR Predator to mouse primary hepatocytes. Taken together, these data provide preliminary proof for the clinical potential of the Predator system.

## 3. Discussion

A comprehensive understanding of cellular processes requires different kinds of tools to identify the function of proteins in the context of cells and organisms(*33*). Genetic methods(*3–6, 20*) and methods targeting RNA level(*7, 14*) have been widely used in researches. However, these methods dependents on the turnover of the target protein for not working at the protein level directly, which can be powerless and problematic when studying long-lived proteins or highly dynamic processes(*1, 8*). Therefore, harnessing the natural UPS to deplete protein targets directly is attracting great interests of researchers(*11*).

The general usage of many current TPD methods that hijack UPS like DeGradFP, AID, and conditional degrons in higher eukaryotes are limited due to their requirement of the prior modification at endogenous protein targets(*9, 20*). In recent years, PROTAC has emerged as a very promising and powerful approach to degrade POI directly both *in vitro* and *in vivo*, however, the time-consuming process for PROTAC design and development, universality, off-target effect and hook effect are major disadvantages(*14, 34–37*). The Trim-Away method is a rapid and acute TPD method, but the high cost of antibody production, the short lifespan of injected-antibody inside the cell, and the technique complexity limit its wide application(*11, 38*).

In the present study, we have developed an innovative and modularized targeted protein degradation system, the Predator system, based on the core function of Trim21(*17, 19, 39*), and we demonstrated that this novel method could directly degrade endogenous target proteins in cells with high efficiency. Compared to Trim-Away technology which introduces antibodies into cells by microinjection or electroporation, our Predator system, through ectopic expression of modularized bifunctional molecules that functions similarly as the antibody to recruit E3 ligase Trim21 and bind to the targets with high specificity, was much more modularized and convenient, because it need only to express the designed Predator system via easy-to-use gene vectors like plasmids or virus (Fig 6)(*9*).

**Figure 6.**
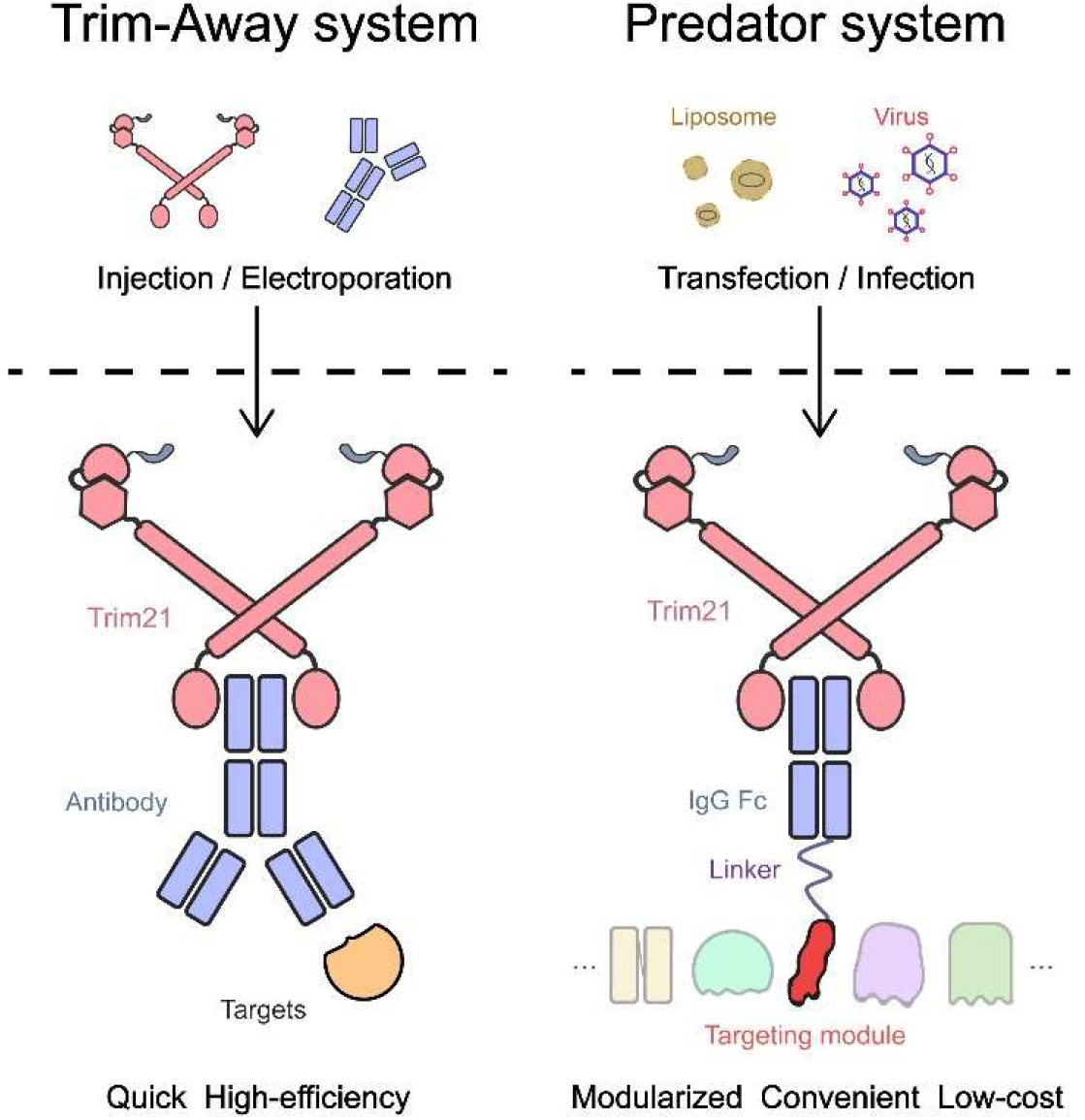
the comparison between Trim-Away and the Predator system.

Owing to the modularized design of the Predator system, both the targeting module and E3 recruitment module are easily modifiable and replaceable. By replacing GFP targeting modules (GFP nanobody) with a single-chain antibody of ErbB3 (ScFv_ErbB3_), we easily changed the orientation of the degradation system and demonstrated that the Predator system can degrade protein targets that are in different physiological states in cells while maintaining high efficiency. These findings suggested that the predator system might have broad application fields, as the substrate specificity of the Predator system is directly dependent on the targeting modules (like nanobody and ScFv we used) that can be adapted to target virtually any protein. Also, there are various kinds of specific interactions to choose for the design of the targeting modules like antibody-antigen interaction, ligands-receptor interaction, and peptide aptamer-targets interaction, etc.

Hence, we introduced ligand-receptor-specific interactions to further broaden the selection of targeting modules of the Predator system. Also, we selected the GCGR, a target of great potential for therapeutic usage(*29–32*), to demonstrate that the clinically-used adenoviral vectors can be used for the delivery and functioning of the Predator system, our results showed that GCGR functioned well in primary hepatocytes. Though preliminary, these results demonstrate the clinical potential of the Predator system. However, *in vivo* experiments are still needed for further demonstration.

Taken together, our work presents a convenient and modularized method for target protein degradation. We believed that in the future the Predator system can play an important role both in fundamental researches and clinical applications.

## 4. Materials and method

### 4.1 Cell Culture

Breast cancer cell lines MDA-MB-231, MCF7, A549, HepG2, Hela and human embryonic kidney cell line HEK293T were purchased from American Type Culture Collection (ATCC, Manassas, VA), and cultured in RPMI 1640 Medium (Hyclone) supplemented with 10% fetal bovine serum (FBS; GIBCO, Gaithersburg, MD, USA) and 100 U/ml penicillin and streptomycin (P/S; Hyclone). Cells were contained in a 5% CO_2_ incubator at 37°C.

### 4.2 Ectopic expression of Predator

Plasmids that express Predator were introduced to cells by lipofecmine3000-mediated transfection according to the manufacturer's directions. the optimized working concentration of plasmid is 2ug/well for the 6-well plate.

### 4.3 RNA Extraction and Quantitative Real-Time PCR

Total RNAs were extracted using the TRIzol agent (Ambion) according to the instruction of the manufacturer. Reverse transcription of RNA and quantitative real-time PCR was performed using the NovoStart^®^ Fast SYBR qPCR SuperMix (Novoprotein) according to the manufacturer's instructions. Quantitative RT-PCR was performed in a Roche 480 real-time PCR system. The 2^−ΔΔCt^ method was used to evaluate the gene expression after normalization for expression of the endogenous controls Hprt1. Each experiment was repeated at least three times.

### 4.4 Western Blotting

Total proteins were extracted with RIPA lysis buffer (Beyotime) with Protease Inhibitor Cocktail (Roche) and Phosphatase Inhibitor Cocktail (Roche). Protein samples were separated in sodium dodecyl sulfate (SDS)-PAGE and transferred to polyvinylidene fluoride (PVDF) filter membranes (Millipore, USA). After blocking in phosphate buffer saline (PBS) containing 0.05% Tween-20 and 5% non-fat milk powder, the membranes were incubated with the following primary antibodies: GFP (Santa Cruz Biotech, 1:500), GCGR (Santa Cruz Biotech, 1:1000), Her3 (ErbB3, Cell Signaling Technology, 1:1000), beta-actin (Santa Cruz Biotech, 1:4000), Secondary antibodies were Horseradish peroxidase (HRP)-conjugated anti-mouse IgG (ZB-2305, ZSGB-Bio, 1:4000) or anti-Rabbit IgG(Fc) (ZB-2301, ZSGB-Bio, 1:4000). Subsequent visualization was detected by Digit imaging system (Thermo, Japan), the gray level of the bands was quantitated by ImageJ software.

### 4.5 In vitro Cell Viability Assays

To perform cell viability assays, cells were counted and plated in the well of 96-well plate (1,500 cells per well) 24h after transfection. the cell viability ability was determined using the Cell Counting Kit-8(CCK8) Assay Kit (Dojindo Corp, Japan) according to the manufacturer's protocol: After the 0/24/48/72 h proliferation of cells, the kit reagent dissolved with RPIM1640 Medium to prepare a 10% working reagent. The original medium was removed and 110 ul working reagent was added to each well. After 2 h incubation in the 37°C incubator. The absorbance was measured at 450 nm to calculate the number of cells, the cell viability assays were performed three independent times.

### 4.6 PAS staining

PAS staining assays were performed according to PAS Cell Staining Kit (Solarbio). In brief, cells were fixed by PAS fixing reagent and rinsed with distilled water. Then the cells were treated with periodic acid for about 15 minutes. Rinse well in distilled water. Cover with Schiff’s reagent for 10-20 minutes. Wash in running tap water for 5-10 minutes. stain with Mayer hematoxylin for 1 minute. Wash in tap water and examine cells microscopically.

### 4.7 Glucose Production Assay

HepG2 cells were plated in 6-well cell plates (1 × 10^6^ cells per well) and allowed to grow for 24 or 48 h in low-glucose DMEM. Cells were washed three times with PBS to remove glucose and starved for 24 or 48 h in 3 ml of glucose production medium (glucose- and phenol red-free DMEM containing the gluconeogenic substrates 20 mM sodium lactate and 2 mM sodium pyruvate) in the absence or presence of 10 nM glucagon. A total of 200 μl of the medium was sampled for the measurement of glucose concentration using a glucose assay kit (SolarBio).

### 4.8 Animals, primary hepatocytes and adenovirus infection

Warm up the perfusion buffer (50 ml HBSS, add 500 ul 7.5% Sodium Bicarbonate solution and 50 ul EDTA solution) and collagenase buffer (10 ml HBSS with Ca2^+^/Mg2^+^, add 100 ul 7.5% Sodium Bicarbonate solution, 20 ul 2.5 M CaCl_2_ and Type II Collagenase to light-medium brown) to 37°C. Insert the needle into the right atrium of the C57BL/6 mouse and start the liver perfusion with perfusion buffer. Cut the portal vein and continue for 3 min. Continue with the collagenase solution for about 3-5 min, stop when the liver becomes transparent and swollen. Dissect the liver, put it in a 10 cm dish, add 250 ul extra collagenase buffers on the liver, and incubate in 37°C incubator for 5 min. Re-suspend hepatocytes with 10 ml of 10% BGS DMEM and filter through a cell strainer. Spin down cells and remove the medium, add 10 ml 10% BGS DMEM, and wash one more time before plating. Plate hepatocytes for 1-2 hours. Replace cell culture plate with DMEM medium (0.1% FBS). Add 30μL adenovirus introduced in each well 2 hours later. After 12 hours’ infection, change the medium to the glucose-free DMEM medium with 10% FBS, add glucagon as stimulation. Incubate cells for 24 hours under the proper condition of 37°C, 5% CO_2_.

### 4.9 Statistical Analysis

Statistical analysis was conducted using the R software. All results were presented as the mean ± standard error of the mean (SEM). Student t-test and One-way ANOVA was performed to compare the differences between treated groups relative to their paired controls. p-values are indicated in the text and figures above the two groups compared and p < 0.05 (denoted by asterisks) was considered as statistically significant.

## Notes

### Competing Interest Statement

The authors have declared no competing interest.

